# The Effect of Glycerol on Microbial Community in Industrial Wastewater Treatment Plant

**DOI:** 10.1101/2023.01.23.525285

**Authors:** M A Prawira Negara, B Jayawardhana, G J W Euverink

**Affiliations:** University of Groningen, Nijenborgh 4, 9747 AG Groningen, The Netherlands; University of Jember, Kampus Tegalboto, Jl. Kalimantan No.37, Krajan Timur, Sumbersari, Kabupaten Jember, Jawa Timur 68121, Indonesia

**Keywords:** Glycerol, Microbial population, Nitrogen removal, Saline Wastewater Treatment Plant

## Abstract

In this paper, we developed and analyzed lab-scale reactors that model the industrial saline wastewater treatment plant (SWWTP) used in North Water SWWTP in Delfzijl, the Netherlands. This industrial wastewater treatment plant is different from a typical municipal wastewater treatment plant, where the wastewater has some recalcitrant chemicals that are hard to degrade and contains a high COD-to-nitrogen ratio and a high concentration of NaCl. The process also differs from other standard industrial wastewater plants where the anaerobic process precedes the aerobic process. The proposed lab-scale reactors are shown to be stable and able to represent the studied industrial SWWTP where glycerol is present in abundance, and there is no similar lab-scale model that has investigated the effect of glycerol on the process. The removal of COD (glycerol) and nitrogen in the system and the changes in the microbial community in both reactors was followed over time. Based on the data, we were able to study the growth of the microbial population that is present in the sludge. The result of the experiment showed that glycerol and ammonia were completely removed, and some nitrate was left in the effluent. At the end of the experiment, we determined that the order Actinomycetales dominated the anaerobic reactor since it is known as the organisms that use glycerol as the carbon source and is quite tolerant with a high salt concentration in the influent. On the other hand, the order Flavobacteriales dominated the aerobic reactor as it is correlated with the ammonia concentration.

## 1. Introduction

Wastewater is a big part of human activities’ excess and needs to be treated before it is environmentally safe to be released into the natural water resources (Henze and Comeau (2008)). If it is released while untreated, it will pollute the water resources and result in hazardous ecological, environmental, as well as economic impacts. Wastewater mainly comes from two major sources: human sewage systems and process waste from industries. In the late nineteenth century, the use of chemicals to treat wastewater was the norm. It was not until the early twentieth century that the biological treatment of wastewater was first introduced, and it has become the basis of wastewater treatment worldwide. The biological treatment of wastewater uses activated sludge, which consists of a high concentration of microorganisms living in microbial communities, mainly consisting of bacteria, protozoa, and archaea. These microbial communities consume organic compounds, ammonia, and phosphate for their metabolism and growth. These compounds are incorporated into the microbial cells, thereby reducing the concentration in the wastewater. The cells are separated from the water by sedimentation of the activated sludge in sedimentation tanks. The clarified wastewater is released safely into natural water resources, such as rivers or the sea.

Although the operation of the treatment process seems to be straightforward, the underlying biological-chemical process is complex and difficult to monitor (Davies (2005)). Consequently, process control relies mainly on the physical process variables that can be measured. There is no monitoring and control of microbial activities. It is already known that the dynamical variation in WWTP is mainly because of the time-varying composition of the microbial communities in the reactors, which is mainly influenced by the variation of influent nutrient composition (Shi et al. (2021)). The influent can vary in the flow rate, chemical composition, pH, and temperature, which in turn affect the population dynamics and metabolic process in the activated sludge.

Different from municipal wastewater, where it is comparatively easy to almost completely remove the pollutants from the wastewater, WWTPs that treat industrial wastewater, which is the subject of this paper, usually deal with recalcitrant chemicals that are hard to degrade by the activated sludge. Such industrial wastewater contains harmful chemicals and has more extreme properties, such as low/high pH or high salt concentration, which directly affect the performance of the activated sludge.

External carbon sources are often fundamental to achieving a high cleaning efficiency in sewage treatment. Glycerol showed significant potential as an external carbon source and energy source for microbial growth in industrial microbiology (da Silva et al. (2009)). Previous studies reported glycerol as a suitable carbon source in wastewater treatment and useful for reducing biomass production. It is also used to remove nitrogen from wastewater during municipal wastewater treatment in two independent activated SBR-type sludge reactors (Smyk and Ignatowicz (2017)). In another study, it was shown that the use of glycerol resulted in good denitrification and phosphorus removal (Salamah and Randall (2020)).

The metabolism of glycerol in microorganisms mainly occurs by two distinct and parallel pathways, oxidative and reductive (Yazdani and Gonzalez (2007)). The members of the Enterobacteriaceae family, such as *Klebsiella, Citrobacter*, and *Clostridia* and some acetic acid bacteria, e.g., *Acetobacter* and *Gluconobacter*, also carry out glycerol degradation using both oxidative and reductive pathways (Gupta et al. (2001)).

In this study, we developed and analyzed lab-scale reactors that model an industrial saline WWTP as used in the North Water SWWTP. Using the developed reactors, we monitored the microbial community in the reactors and investigated the effect of glycerol in the influent of this WWTP and how it will influence the removal of nitrogen in the system. The use of lab-scale reactors enables us to gain insight into the microbial activities in WWTP processes. Having the knowledge of microbial activities is important as it will help us in gaining a better picture of the dynamic model of the system. We can use the microbial community dynamics and couple it with the parameter-varying of the dynamic model (see cite simulation paper).

The rest of the paper is organized as follows. In Section 2, we explain the materials and methodology that we use in this study. In Section 3, we present the result derived from the laboratory experiment and the analysis of the resulting data. The paper is then concluded in Section 4.

## 2. Materials and methods

### 2.1. North Water’s wastewater treatment plant

In this study, we used the system that is used by North Water’s Saline Wastewater Treatment Plant (SWWTP) in Delfzijl, the Netherlands. This industrial SWWTP^1^ treats collective industrial wastewater from chemical industries that are operated in a chemical park in Delfzijl and its neighborhood. The processed clean water is then released into the surface water. General operational data of this SWWTP are given in Table 1.

**Table 1.**
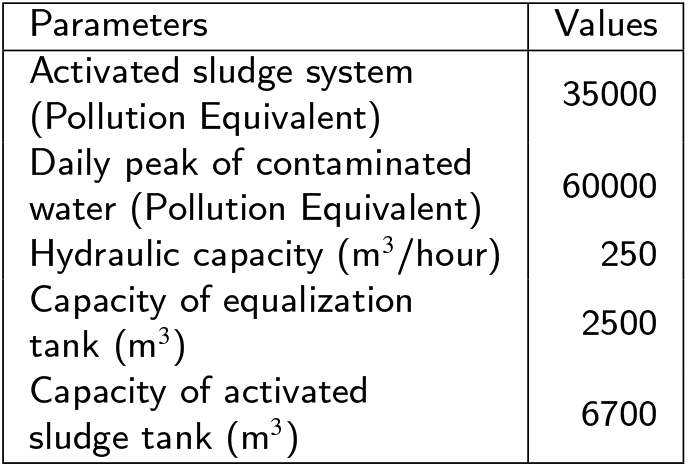
North Water’s SWWTP Purification Data

| Parameters | Values |
| --- | --- |
| Activated sludge system (Pollution Equivalent) | 35000 |
| Daily peak of contaminated water (Pollution Equivalent) | 60000 |
| Hydraulic capacity (m<sup>3</sup>/hour) | 250 |
| Capacity of equalization tank (m<sup>3</sup>) | 2500 |
| Capacity of activated sludge tank (m<sup>3</sup>) | 6700 |

Table 2 shows the average value of the influent for North Water’s SWWTP. These average values are based on the measurements that were performed in week 40 in 2014 until week 10 in 2017, which can be seen in Appendix A.

**Table 2.** Average composition of the influent at North Water’s SWWTP

| Component | Concentration (mg/L) |
| --- | --- |
| BOD | 1000 |
| COD | 2000 |
| Cl | 12000 |
| K | 24 |
| N | 103 |
| Na | 7000 |
| NH<sub>4</sub> | 23 |
| Nkj | 21 |
| NO<sub>2</sub> | 5 |
| NO<sub>3</sub> | 66 |
| PO<sub>4</sub> | 8 |
| SO<sub>4</sub> | 714 |
| TOD | 2200 |
| Glycerol | 1600 |
| Methanol | 165 |

A general schematic of the biological treatment process of North Water SWWTP is shown in Fig. 1. The influent coming from various sources is collected in the influent tank before it is pumped into the inner part of a 2-zone treatment tank where an anaerobic biological process takes place and an aerobic process in the outer part of the treatment tank. In the clarifier tank, the activated sludge is separated from the water and is partly fed back into the treatment tank, while the treated water is then discharged into the Wadden Sea.

**Figure 1:**
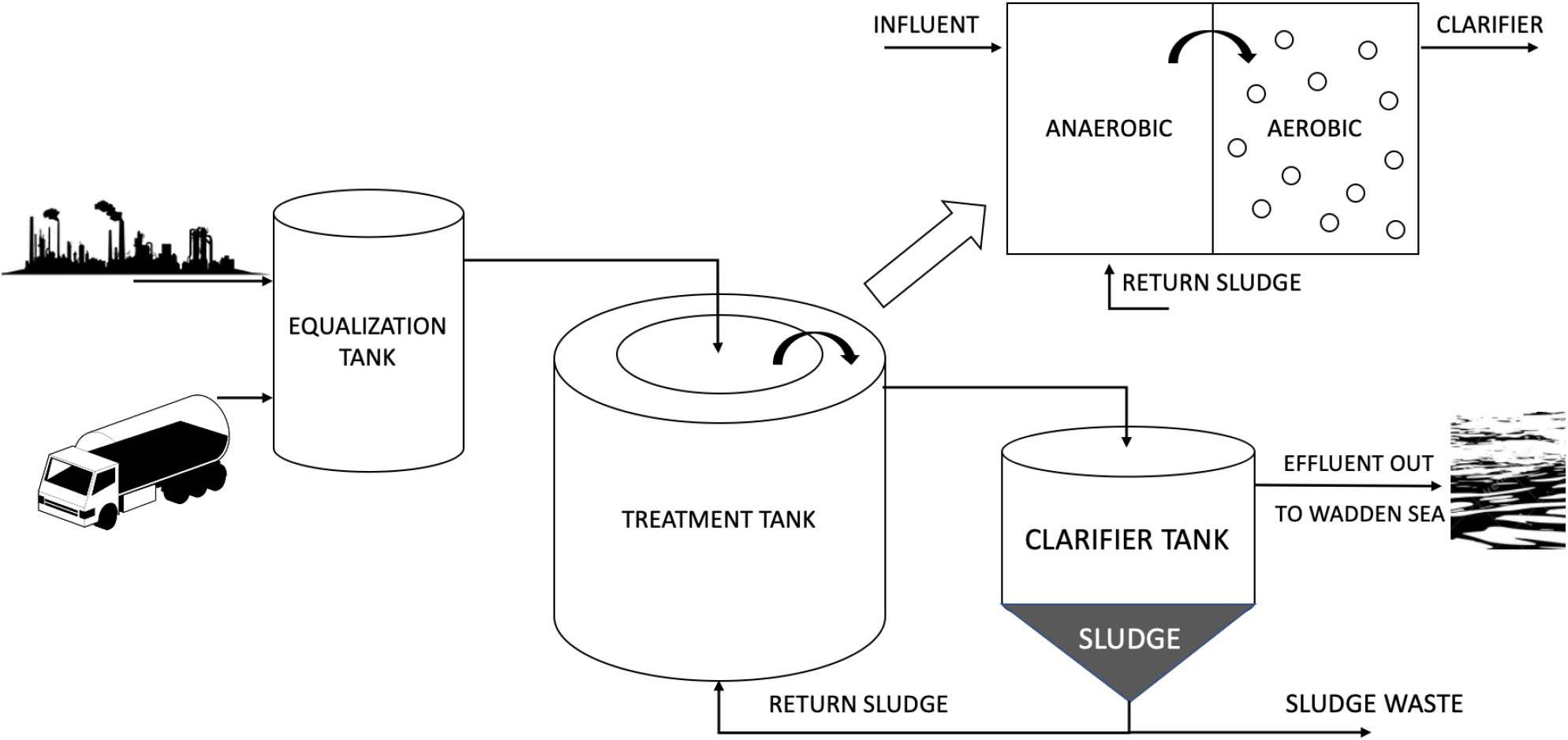
Schematic diagram of the North Water’s Saline Wastewater Treatment Plant.

The setup of an anaerobic process followed by an aerobic one is not common for a standard WWTP. This reverse setup is possible in North Water’s WWTP since the influent has a high chemical oxygen demand (COD) such that it contains sufficient energy in the wastewater for the biological removal process of 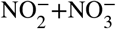 in the first reactor. The further removal of contamin^2^ ants ta^3^kes place in the outer reactor part, where industrial waste is further broken down by biological processes.

### 2.2. Reactors set-up

We conducted our experiment based on North Water’s SWWTP, which resulted in the experimental setup using a two-reactor, as can be seen in Fig. 2, where the first reactor is anaerobic and the second reactor is aerobic. We also separate the influent into two different vessels (Carbon and Nitrogen) to avoid too much growth in the influent vessels. As shown in Fig. 2, the influent coming from the two vessels is pumped to the first reactor, where the anaerobic biological process takes place before it is then pumped into the second tank, where the aerobic process occurs. After that, it is then pumped into the effluent vessel. There is a slightly different approach from North Water’s SWWTP operation, where we do not add return sludge from the effluent back into the first reactor.

**Figure 2:**
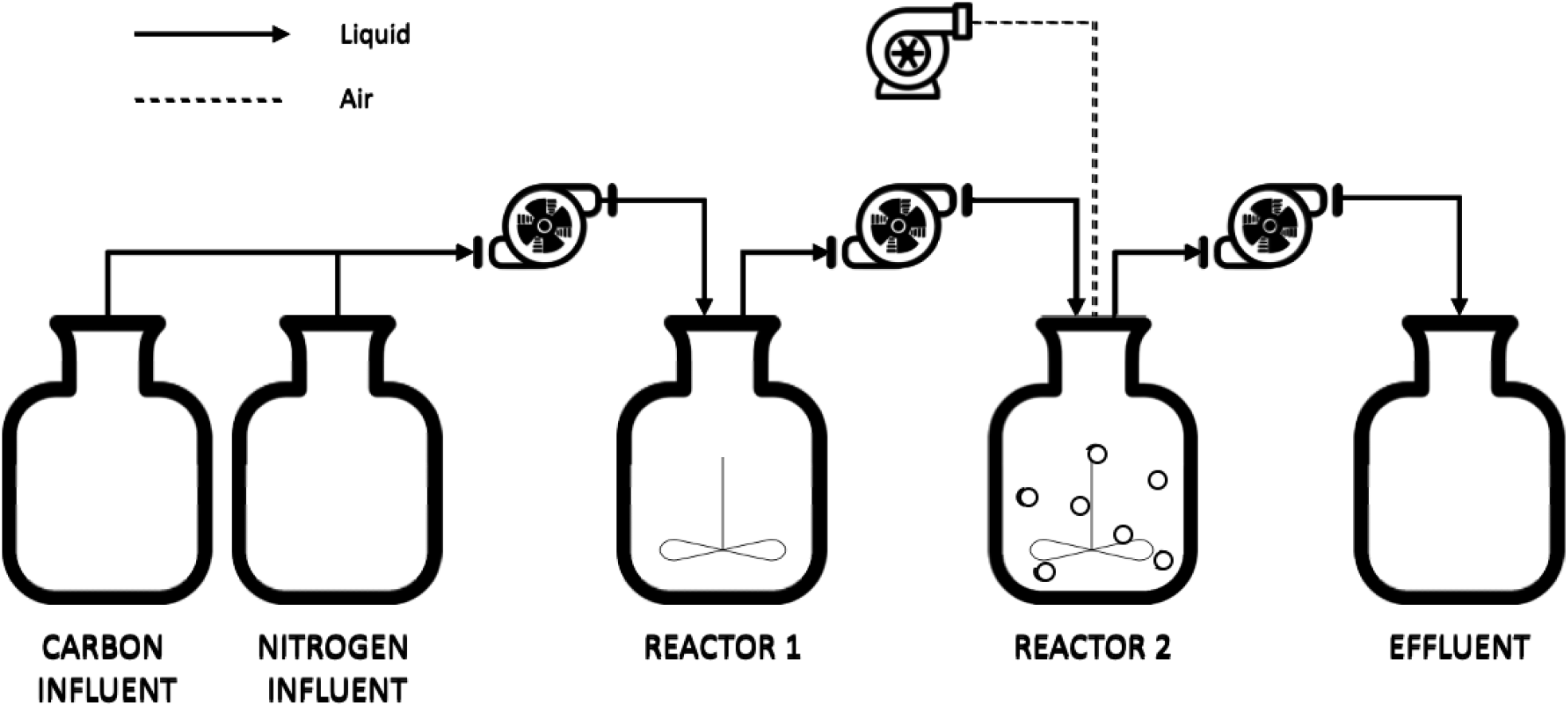
Schematic diagram of the experiment setup based on North Water’s SWWTP.

### 2.3. Experimental start-up and operation

The experiment was divided into three sequential periods that ran continuously. The first part was the initializing part, where we used the sludge and influent from North Water to start-up the reactors in the laboratory. We ran the system for four weeks, after which the influent was replaced with a synthetic influent in the second period. After eight weeks, the composition of the synthetic influent was changed. Samples were taken at regular intervals to determine the COD, ammonia, nitrate, and microbial composition.

The composition of the synthetic wastewater that was used throughout the experimental period can be seen in Table 3. As shown in Fig. 2, we separated the influent into two different solutions, carbon influent and nitrogen influent. The carbon influent is a solution of glycerol, methanol, and magnesium sulfate, while the rest of the components are in the nitrogen influent.

**Table 3.** Synthetic influent’s composition

| Influent Solution | Component | Concentration (mg/L) |
| --- | --- | --- |
| Carbon Influent | Glycerol | 4000 |
| Carbon Influent | Methanol | 160 |
| Nitrogen Influent | (NH<sub>4</sub>)<sub>2</sub>SO<sub>4</sub> | 300 |
| Nitrogen Influent | NaCl | 20240 |
| Nitrogen Influent | KNO<sub>3</sub> | 100 |
| Nitrogen Influent | KH<sub>2</sub>PO<sub>4</sub> | 182 |
| Nitrogen Influent | K<sub>2</sub>HPO<sub>4</sub> 3H<sub>2</sub>O | 278 |
| Carbon Influent | MgSO<sub>4</sub> | 100 |
| Nitrogen Influent | Trace Element | 1 |

### 2.4. Chemical Analysis

The chemical composition and the pH of both reactors were determined twice a week. The COD was measured using LCK314, the ammonia was measured using LCK303, and the nitrate was measured using LCK339 (Hach, German).

### 2.5. High-throughput 16S-rDNA gene sequencing and analysis

The frozen samples (−20°C) were thawed at room temperature and total DNA was extracted using the FastDNA® Spin Kit for Soil (MP Biomedicals, USA) according to the manufacturer’s protocol. The extracted genomic DNA was used as the template in the PCR reactions. Part of the 16S-rDNA genes was amplified using the primers 338F (‘5-ACT CCT ACG GGA GGC AG) and 805R (‘5-GAC TAC CAG GGT ATC TAA TCC). We also multiplexed the amplified DNA barcoded prior to the sequencing procedure using Illumina Nexseq 1000. Consequently, we obtained the diversity profiles by plotting the number of 16S-rDNA sequences as a percentage in a stacked bar chart.

## 3. Result and discussion

### 3.1. pH, COD, and Nitrogen

The result of the measurements that were taken for around eleven weeks, starting from 06/07/2021 until 16/09/2021, are shown in Fig. 3. The soluble COD decreased in R1 to approximately 50% of the influent and reduced further to less than 100 mg/l in R2. Ammonium is removed in R1 to 20% of the initial concentration in the influent during the whole experiment. In R2, most of the ammonia is further removed to below the detection limit in the first phase. In the second phase, when the COD content in the influent was increased by 20%, not all ammonia was removed in R2, and no significant changes in ammonia removal were observed in R1. Some acids were produced in R1, as can be concluded from the decrease in pH. In R2, the acids are consumed by the microorganisms, and the pH increases to the initial pH of the influent.

**Figure 3:**
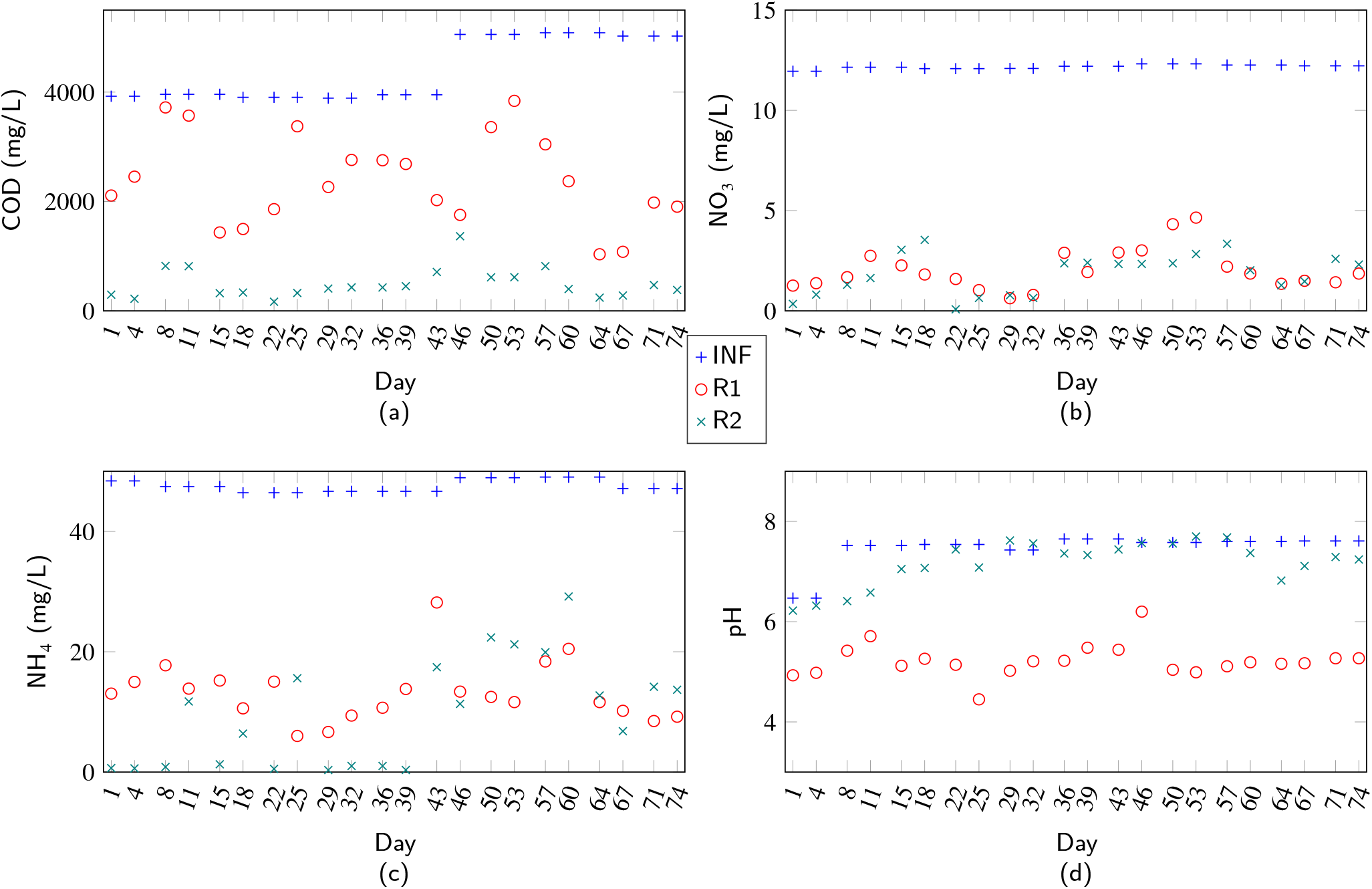
Lab measurement results for chemical composition and acidity are taken from 05/07 until 16/09. (a) COD (b) Nitrate Ammonia (d) acidity

In R1, more than 90% of the incoming nitrate was removed in R1 only. No further reduction was observed in R2. After the increase of the COD in the second phase, no changes in nitrate removal were determined. In the anaerobic R1, most likely, nitrate serves as the e-acceptor. In the aerated R2, oxygen became the preferred electron acceptor and the concentration of nitrate did not decrease further (Fig. 3 (b)).

### 3.2. Microbial Community

The composition of the microbial community in the two-reactor setup, R1 and R2, was analyzed using NGS of the 467 bp PCR product. DNA was isolated from bacteria that were present in the reactors at different time points during the feeding with the artificial wastewater. The PCR products were barcoded, pooled, and sequenced on the NEXTgen 1000. In Table 4, the general characteristics of the sequencing are shown. The sequences were used to identify which taxonomic order and their relative abundance were present in the reactors at the different time points. The sequencing data are shown in Fig. 4.

**Table 4.** General characteristics of the sequencing data

| Sample | Reactor | Number of reads | Number of unique sequences |
| --- | --- | --- | --- |
| 1 | 1 | 109190 | 18779 |
| 2 | 1 | 99348 | 6041 |
| 3 | 1 | 387191 | 48127 |
| 4 | 1 | 90447 | 11207 |
| 1 | 2 | 78067 | 21721 |
| 2 | 2 | 95595 | 14921 |
| 3 | 2 | 326017 | 103960 |
| 4 | 2 | 92509 | 15624 |

**Figure 4:**
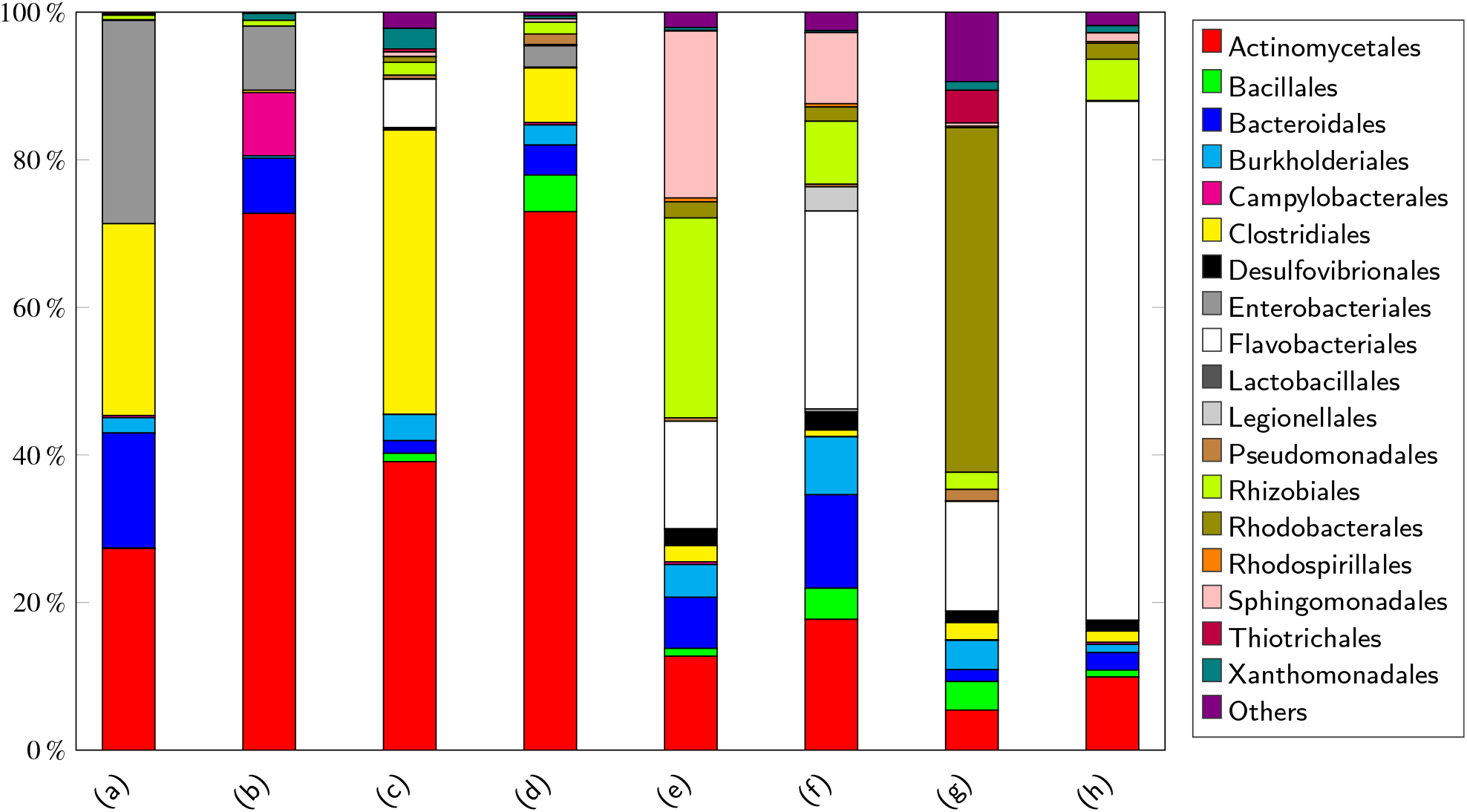
Stacked-bar-chart of microbial communities showing the population in order for few samples, where (a), (b), (c) and are R1 samples while (e), (f), (g) and (h) are R2 samples. Samples (a) and (e) are taken on day 8, samples (b) and (f) are taken on day 15, samples (c) and (g) are taken on day 43, and samples (d) and (h) are taken on day 53.

The top 5 of the bacteria in Reactor 1 contained members of the order Actinomycetales, Enterobacteriales, Clostridiales, Bacteriodales, and Burkholderiales. Reactor 1 was run at anoxic conditions, and the top 5 of the order all contain members that can grow under oxygen limitation or at anaerobic conditions. As shown in Fig. 4, in the second sample of R1 the Actinomycetales increased to 70% of the population, and the Clostridiales and Burkholderiales were not detected. At this time point, 10% of the population were members of the Campylobacteriales. Actinomycetales remained the largest part of the population throughout the experiment, and the Clostridiales reappeared in R1 in the third and fourth samples.

In the artificial wastewater, the major carbon source was glycerol (4 g/l) and a minor portion of methanol (0.16 g/l). Several species of the detected order are able to metabolize glycerol and methanol under oxygen-limited conditions. E.g., *E. coli, Citrobacter freundii*, and *Klebsiella pneumoniae* from the Enterobacteriales and also *Clostridium pasteurianum* and *Clostridium butyricum* from the Clostridiales were previously shown to use glycerol as carbon source (Clomburg and Gonzalez (2013)). In the aerobic reactor, R2 also contains microorganisms that are able to degrade, e.g., *Streptomyces natalensis* from Actinomycetales was previously shown to use glycerol as a carbon source (Recio et al. (2006)). In other research works from Kaiser et al. (1994) and Borodina et al. (2005), it is concluded that glycerol can be a good carbon source for the other Streptomyces species. Some species of Bacillales and Rhodobacterales, e.g., *Bacillus methanolicus* and *Paracoccus denitrificans*, are able to consume methanol (see Verseveld and Stouthamer (1978) and Krog et al. (2013)). The microbial consortium is apparently not able to degrade all of the COD at the dilution rate and anoxic conditions that were used for R1 (see Fig. 3 (a)). The effluent of R1 is passed to R2. The volume of R1 and R2 were the same, so the dilution rate for both reactors was the same.

The top 5 bacteria in R2 contained members of the order Flavobacteriales, Rhodobacterales, Rhizobiales, Actinomycetales, and Sphingomonadales. Because the presence of oxygen is higher in R2 than in R1, the COD can be further decomposed in the anaerobic reactor, see Fig. 3 (a). Looking at Fig. 4, we can see that in the population, the Rhizobiales decreased from the second sample until the last sample. The opposite occurred with the Flavobacteriales, and they became the major part of the microbial community in R2 in the last sample.

We can explain that the high presence of Actinomycetales in both R1 and R2 is because Actinomycetales can tolerate the high salt concentration in the influent (we have around 3% salt concentration in our media). Other salt-tolerant representatives of the order are also present in the two reactors, e.g., Bacillales, Enterobacteriales, Pseudomonadales, Lactobacillales, and Sphingomonadales (see Fatima and Arora (2019) and Kearl et al. (2019)).

Another thing that we can gain from the sequencing result of both reactors is when we relate it to the lab measurement results that are shown in Fig. 3 (d), we can see that for pH reading that shows the value of reactor 1 to be lower than reactor 2. We can relate this result to the higher combined population of fermentative bacteria in Reactor 1 compared to Reactor 2, such as Enterobacteriales, Clostridiales, and Lactobacillales. Furthermore, if we focus more on the lab measurement result on the same date as the sequencing sample, which can be seen in Fig. 3, we can also see a few more relations between the lab measurement result and the sequencing result.

The microbial community in R1 consists of a fair amount of fermentative microorganisms, e.g., Enterobacteriales, Clostridiales, and Lactobacillales. This is in agreement with the increase in acidity to pH 5 on average. The R1 at days 8 and 43, as shown in Fig. 3 (c) is higher than at days 15 and 53. The reason might be the increased relative abundance of Clostridiales and Enterobacteriales (Fig. 4). According to Caskey and Tiedje (1980), *Clostridium sp*. from Clostridiales can produce ammonia from protein and amino acids. Feeding these organisms to the decaying cells in the reactor will produce ammonium. Likewise, based on the work of Stewart (1994), *E. coli* and *Klebsiella pneumoniae* the Enterobacteriales also can produce ammonia amino acids.

Fig. 3 (c) also shows that at day 8, the concentration of nitrate and ammonia in the R2 is the lowest compared to days 15, 43, and 53. This happened because the concentration of Rhizobiales is the highest, just as shown in Fig. 4. According to Kutvonen et al. (2015), *Rhizobium, Mesorhizobium*, and *Tardiphaga* from Rhizobiales used ammonium and nitrate as nitrogen sources. Another thing that we can see in Fig. 3 (c) is the value of ammonia in the R2 at day 53, which shows the highest value compared to days 8, 15, and 43. This happened because the concentration of Flavobacteriales is the highest, just as shown in Fig. 4. Based on the work of Muck et al. (2019), the order Flavobacteriales was positively correlated with ammonia concentration.

## 4. Conclusions

The proposed lab-scale SWWTP reactors have been shown to be stable and can be used to model the biological process in industrial SWWTP. It has been used to show that glycerol can be used as the carbon source for the reactors and is able to help in the removal of nitrogen by the order of Rhizobiales and Flavobacteriales, among others, which consume glycerol as carbon source. The lab-scale reactors were able to remove glycerol and ammonia completely with low amount of nitrate in the effluent. Analysing the dynamics of microbial population in the two reactors R1 and R2, the R1 consists of a fair amount of fermentative microorganisms, which is shown in the acidity value of pH 5 on average. In R1, the concentration of ammonia might be related to the amount of Clostridiales and Enterobacteriales. On the other hand, the abundance of Rhizobiales and Flavobacteriales in R2 affected the concentration of ammonia and nitrate. At the end of the experiment, we can also see that the order Actinomycetales dominates the anaerobic reactor while the order Flavobacteriales dominates the aerobic reactor.

Further work on this research is to use the knowledge of the microbial activities and also the chemical composition in the two reactors to create a dynamic model of the system, such as using a well-known Activated Sludge Model and coupling some of the microbial population to some of the parameters to get a parameter-time-varying based on the microbial dynamics.

## Acknowledgement

M A Prawira Negara profoundly expresses gratitude to the Islamic Development Bank (IsDB) for the scholarship funding.

